# Kinase activity of TBK1 is required for its binding to STING, but not for its recruitment to the Golgi

**DOI:** 10.1101/2021.04.27.441586

**Authors:** Haruka Kemmoku, Yoshihiko Kuchitsu, Kojiro Mukai, Tomohiko Taguchi

**Author notes:** Correspondence and requests for materials should be addressed to K. Mukai and T. Taguchi.

## Abstract

Stimulator of interferon genes (STING) is an innate immune protein for DNA pathogens. In response to the emergence of DNA in the cytosol, STING relocates from the endoplasmic reticulum (ER) to the Golgi and induces the type I interferon response through cytosolic TANK-binding kinase 1 (TBK1). The molecular mechanism underlying TBK1 activation by STING remains poorly understood. Here we report a cell system by which STING and TBK1 are simultaneously monitored. The system utilizes STING/TBK1-double knockout (KO) cells, fluorescent protein-tagged TBK1 and STING, and super-resolution microscopy. After STING stimulation, TBK1 is directly recruited to the trans-Golgi network (TGN), not to the other parts of the Golgi. The recruitment of TBK1 does not require its kinase activity. C-terminal STING variants (ΔC9 and L373A), in which the TBK1-STING binding interface is mutated or deleted, induce the recruitment of TBK1. These results indicate that the kinase activity of TBK1 or the C-terminal motif of STING is not required for its recruitment to TGN, but rather for the formation of the stable STING signalling complex at TGN.

## Introduction

The detection of microbial pathogens with nucleic acid sensors is a central strategy of innate immunity (Palm and Medzhitov, 2009; Takeuchi and Akira, 2010). The nucleic acid sensors localize at various subcellular compartments and, upon binding to foreign nucleic acids, trigger potent immune responses for host defense (Wu and Chen, 2014). Cyclic GMP-AMP synthase (cGAS) is a sensor for double-stranded DNA in the cytosol (Sun et al., 2013). It synthesizes cyclic GMP-AMP (cGAMP) with ATP and GTP (Wu et al., 2013), which stimulates the induction of type I interferons and proinflammatory cytokines through the cGAMP sensor STING (Ishikawa and Barber, 2008) [also known as MITA (Zhong et al., 2008), ERIS (Sun et al., 2009), MPYS (Jin et al., 2008), or TMEM173]. STING is postulated to act as a scaffold to activate the downstream protein kinase TBK1 (Tanaka and Chen, 2012; Zhang et al., 2019). Activated TBK1 phosphorylates and thus activates IRF3, the essential transcription factor that induces the synthesis of type I interferons (Fitzgerald et al., 2003). During this process, TBK1 phosphorylates also STING, generating the IRF3-docking site on STING (Liu et al., 2015).

STING is an ER-localized transmembrane protein (Ishikawa and Barber, 2008). After STING binding to cGAMP, STING exits the ER with COP-II vesicles. The COP-II-mediated translocation of STING from the ER is required to activate the downstream STING signalling pathway (Dobbs et al., 2015; Mukai et al., 2016; Ogawa et al., 2018; Gui et al., 2019; Sun et al., 2018; Ran et al., 2019). Phosphorylated TBK1, the active form of TBK1, is exclusively localized to a subdomain within the TGN (Mukai et al., 2016). Palmitoylation of STING that occurs at the Golgi is essential to activate the downstream signalling pathway (Mukai et al., 2016; Hansen et al., 2018; Haag et al., 2018; Jia et al., 2020). In two autoinflammatory monogenic diseases, STING-associated vasculopathy with onset in infancy (SAVI) (Liu et al., 2014) and the COPA syndrome (Watkin et al., 2015), STING is constitutively activated without DNA stimulation and localizes not to the ER but to the perinuclear compartments including the Golgi (Jeremiah et al., 2014; Mukai et al., 2016; Ogawa et al., 2018; Lepelley et al., 2020; Deng et al., 2020; Mukai et al., 2021). Together, these results demonstrate that the Golgi is an organelle at which STING activates TBK1 for triggering the STING-dependent innate immune response (Taguchi and Mukai, 2019). However, the precise mechanism underlying the specific activation of TBK1 at the Golgi, not the ER, remains unclear.

Recent studies show that a conserved PLPLR[T/S]D motif within the C-terminal tail of STING mediates the activation of TBK1 (Zhang et al., 2019; Zhao et al., 2019). In this work, we develop a cell system by which STING and TBK1 are simultaneously monitored. With this system, we examined the contribution of the PLPLR[T/S]D motif of STING and the kinase activity of TBK1 to the recruitment of TBK1 to the Golgi.

## Results

### Generation and validation of cells with fluorescent protein-tagged TBK1 and STING

To monitor dynamics of TBK1 and STING, we tagged TBK1 and STING with mScarletI and mNeonGreen, respectively. Mouse embryonic fibroblasts (MEFs) were chosen since the cGAS/STING pathway is active in these cells. We prepared STING/TBK1 double knockout MEFs

(ST-DKO MEFs) from STING knockout MEFs using the CRISPR-Cas9 technology to eliminate the contribution of endogenous STING and TBK1. ST-DKO MEFs were then reconstituted with *C*-terminally mScarletI-tagged TBK1 and *N*-terminally mNeonGreen-tagged STING.

We then validated if fluorescent protein-tagged STING and TBK1 are functional. Cells were stimulated with DMXAA, a membrane-permeable mouse-specific STING agonist (Prantner et al., 2012). As expected, 60 min after DMXAA stimulation, we observed phosphorylated TBK1 at Ser172 (p-TBK1), phosphorylated STING at Ser365 (p-STING), and phosphorylated IRF3 at Ser396 (p-IRF3) in wild-type (WT) MEFs, but not in ST-DKO MEFs (Figure 1A). In ST-DKO MEFs reconstituted with mNeonGreen-STING and TBK1-mScarletI, phosphorylated TBK1-mScarletI, phosphorylated mNeonGreen-STING, and p-IRF3 were detected after DMXAA stimulation, showing that reconstituted two proteins were functional.

**Figure 1.**
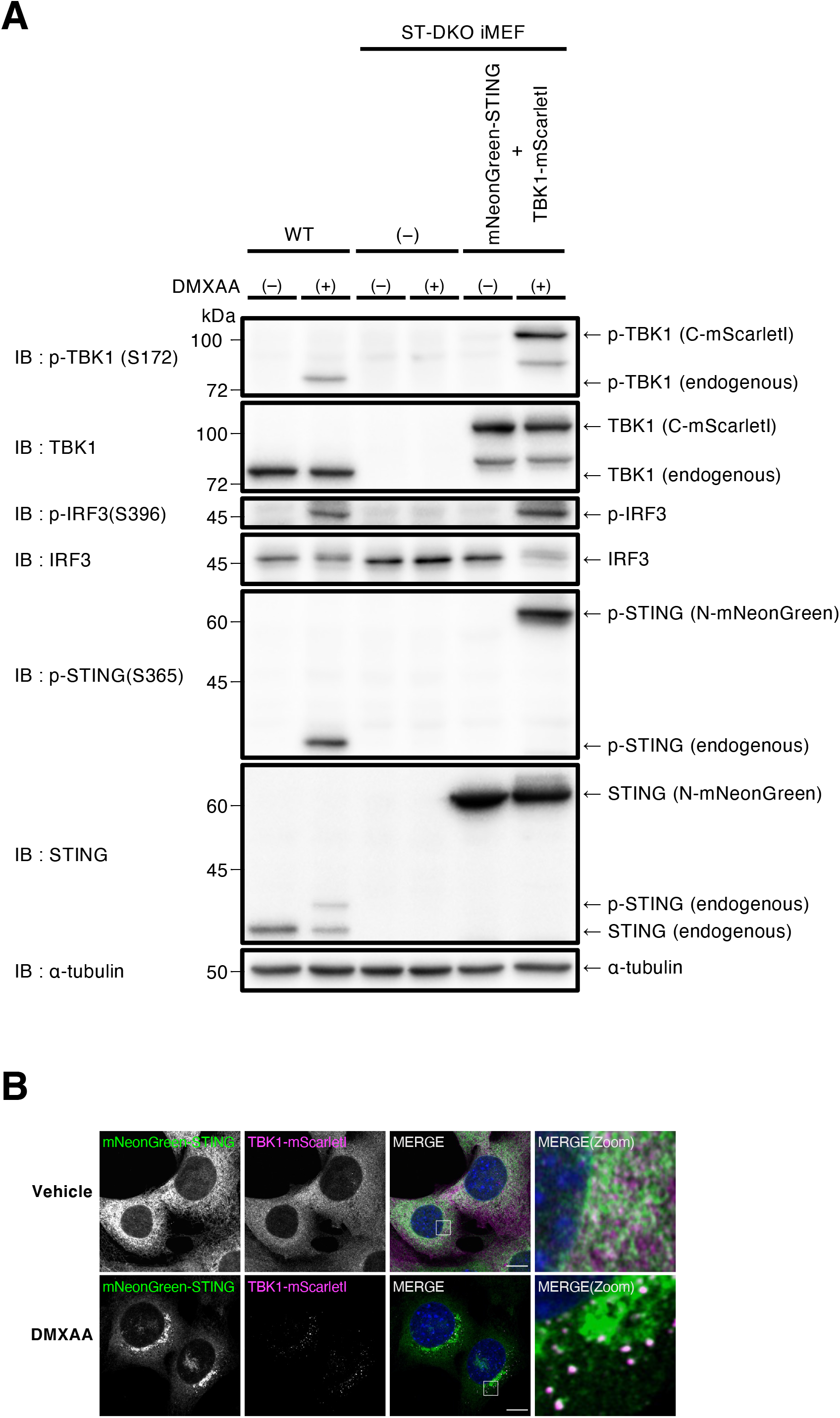
Generation of mNeonGreen-STING- and TBK1-mScarletI-reconstituted MEFs. (A) STING/TBK1 double-knockout MEFs (ST-DKO MEFs) were generated from STING-knockout MEFs using the CRISPR-Cas9 technology. ST-DKO MEFs were reconstituted with mNeonGreen-STING and TBK1-mScarletI. The reconstituted ST-DKO MEFs were stimulated with DMXAA (25 μg/mL) for 60 min. Cell lysates were prepared and analyzed by western blot. (B) mNeonGreen-STING- and TBK1-mScarletI-expressing ST-DKO MEFs were stimulated with DMXAA (25 μg/mL) for 60 min, fixed, permeabilized, and stained with DAPI (blue). Scale bar, 10 μm.

As with endogenous STING (Konno et al., 2013) and TBK1 (Wang et al., 2014; Ogawa et al., 2018) in unstimulated cells, mNeonGreen-STING and TBK1-mScarletI showed typical ER and cytosolic localization, respectively (Figure 1B). After stimulation, STING exits the ER and is transported to the perinuclear compartments including the Golgi (Mukai et al., 2016). mNeonGreen-STING also relocated to perinuclear region, 60 min after stimulation with DMXAA. At this time point, TBK1-mScarletI drastically changed its localization. TBK1-mScarletI relocated from the cytosol to multiple puncta, at which TBK1-mScarletI co-localized with mNeonGreen-STING.

### TBK1 is recruited to TGN, not to the other parts of the Golgi

The Golgi is a polarized organelle that has distinct functional domains, such as the cis-Golgi network (CGN) and TGN. Treatment of cells with a microtubule-depolymerizing agent nocodazole results in dispersed Golgi stacks “mini-Golgi” in the cytoplasm, and this fragmentation facilitates the analysis of *cis*-to-*trans* polarity of the Golgi (Shima et al., 1997). We treated cells with nocodazole for 90 min and then with DMXAA for 30 min. As expected, GM130 (a CGN protein) and TGN38 (a TGN protein) were scattered throughout the cytoplasm (Figure 2A). One mini-Golgi indicated by arrowhead (Figure 2A) was magnified and analyzed (Figures 2B and 2C). Within the mini-Golgi, while mNeonGreen-STING localized at both CGN and TGN, TBK1-mScarletI was confined to TGN. Analysis of multiple mini-Golgis also showed the exclusive confinement of TBK1-mScarletI to sub-domains within TGN (Figure S1). Activated TBK1 phosphorylates STING on Ser365 (Liu et al., 2015). The phosphorylated STING was also confined to sub-domains within TGN (Figure S2). These results suggested that TBK1 was directly recruited to TGN from the cytosol, but not to the other parts of the Golgi and was activated at TGN.

**Figure 2.**
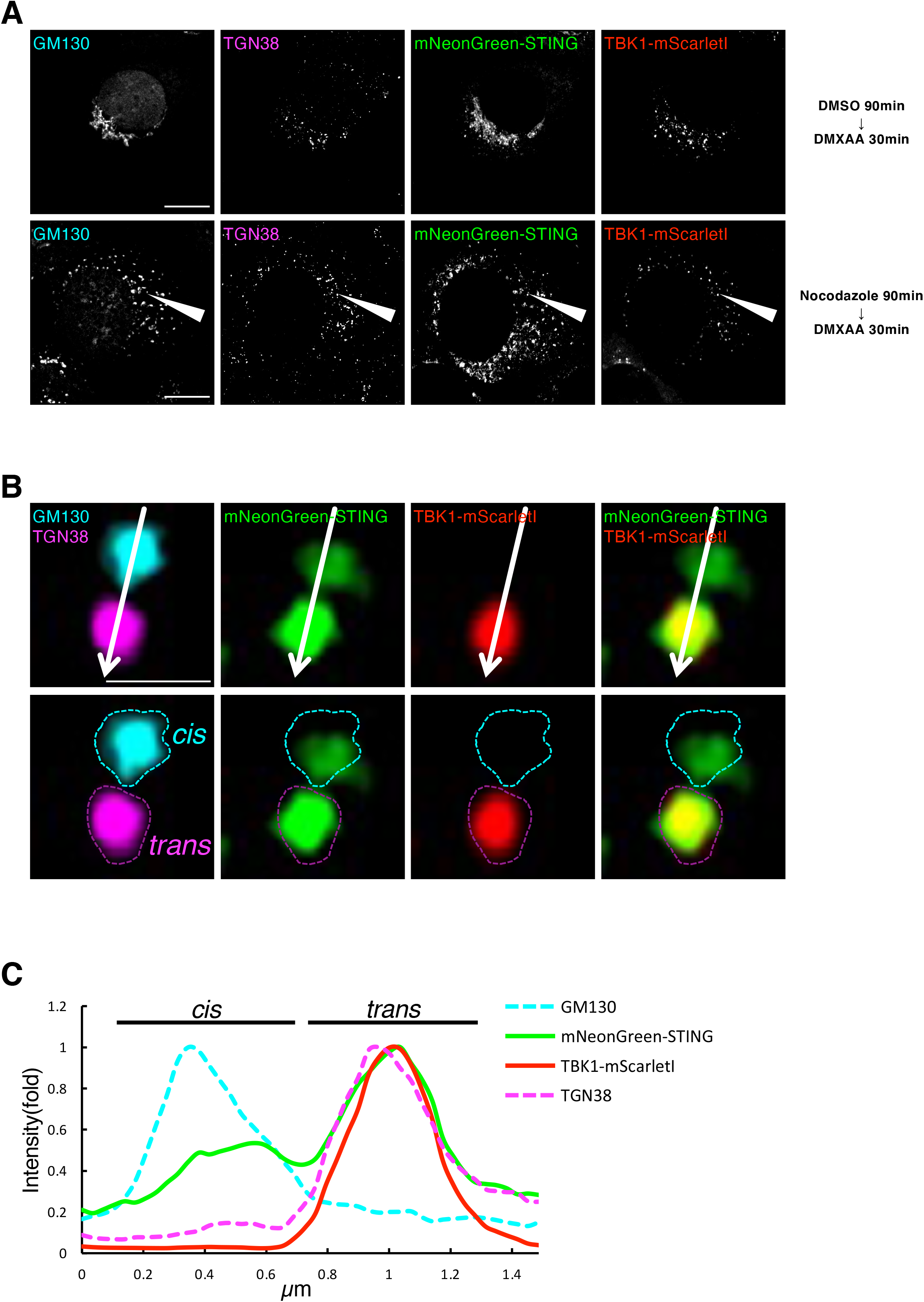
TBK1 is recruited to TGN, not to the other parts of the Golgi. (A) mNeonGreen-STING- and TBK1-mScarletI-reconstituted ST-DKO MEFs were treated with nocodazole (2.5 μM) for 90 min, followed by stimulation with DMXAA (25 μg/mL) for 30 min. Cells were fixed, permeabilized, and stained for GM130 (a CGN protein, cyan) and TGN38 (a TGN protein, magenta). Scale bars, 10 μm. (B) One mini-Golgi indicated by arrowhead in (A) was magnified. The *cis*- and *trans*-regions of the mini-Golgi were outlined in the images at the bottom row. Scale bar, 1 μm. (C) Fluorescence intensity profile along the arrows in (B) is shown. See also Figure S1.

We sought to examine the site of TBK1 recruitment to intact Golgi. To this end, we resorted to use of Airyscan super-resolution microscopy. Airyscan is a 32-channel gallium arsenide phosphide photomultiplier tube (GaAsP-PMT) area detector that collects a pinhole-plane image at every scan position, providing lateral resolution to even 120 nm for two-dimensional imaging (Huff et al., 2017). To visualize CGN or TGN, *N*-terminally mScarletI-tagged GM130 or Rab6 (a TGN protein) was stably expressed in TBK1-knockout MEFs that were reconstituted with TBK1-mNeonGreen. With by Airyscan superresolution microscopy, CGN and TGN could be resolved, although not completely, as indicated by different localization profiles between mScarletI-GM130 (CGN) and TGN38 (TGN) or between GM130 (CGN) and mScarletI-Rab6 (TGN) (Figure S3). Cells were stimulated with DMXAA for 20 min, fixed, and analyzed by Airyscan superresolution microscopy. TBK1-mNeonGreen formed multiple perinuclear puncta after DMXAA stimulation (Figures S4A-S4H) and individual puncta were examined if they were within CGN or TGN. The number of TBK1 puncta on TGN was significantly larger than that on CGN (Figures S4I), reinforcing that TBK1 was directly recruited to TGN.

### PLPLR[T/S]D motif of STING is dispensable for TBK1 recruitment

Recent structural studies indicate that the PLPLR[T/S]D motif within the C-terminal tail of STING binds the dimer interface of TBK1 (Zhang et al., 2019; Zhao et al., 2019). Mutations or deletion of amino acid residues in the motif suppress the activation of TBK1, phosphorylation of STING, and the induction of type I interferons (Zhang et al., 2019; Zhao et al., 2019). We thus examined if STING variants with mutation or deletion in the motif can recruit TKB1 to the Golgi. We chose STING (ΔC9) in which the nine C-terminal residues (^364^<u>PLPLRTD</u>LI) is deleted, and STING (L373A), because these two STING variants mostly lose the ability to induce the type I interferons (Zhao et al., 2019).

*N*-terminally FLAG-tagged STING variants [FLAG-STING (ΔC9) or FLAG-STING (L373A)] and TBK1-mScarletI were reconstituted in ST-DKO MEFs. After stimulation with DMXAA for 60 min, p-TBK1 (Ser172), p-STING (Ser365), and p-IRF3 (Ser396) emerged in MEFs expressing wild-type (WT) STING, but not in MEFs expressing STING (ΔC9) or STING (L373A) (Figure 3A). After stimulation, the amount of TBK1 that was co-immunoprecipitated with FLAG-STING increased about 5-fold in MEFs expressing WT-STING, but did not increse in MEFs expressing STING (ΔC9) or STING (L373A). These results were consistent with the previous findings that the PLPLR[T/S]D motif within the C-terminal tail of STING is essential for the binding to and activation of TBK1 (Zhao et al., 2019).

**Figure 3.**
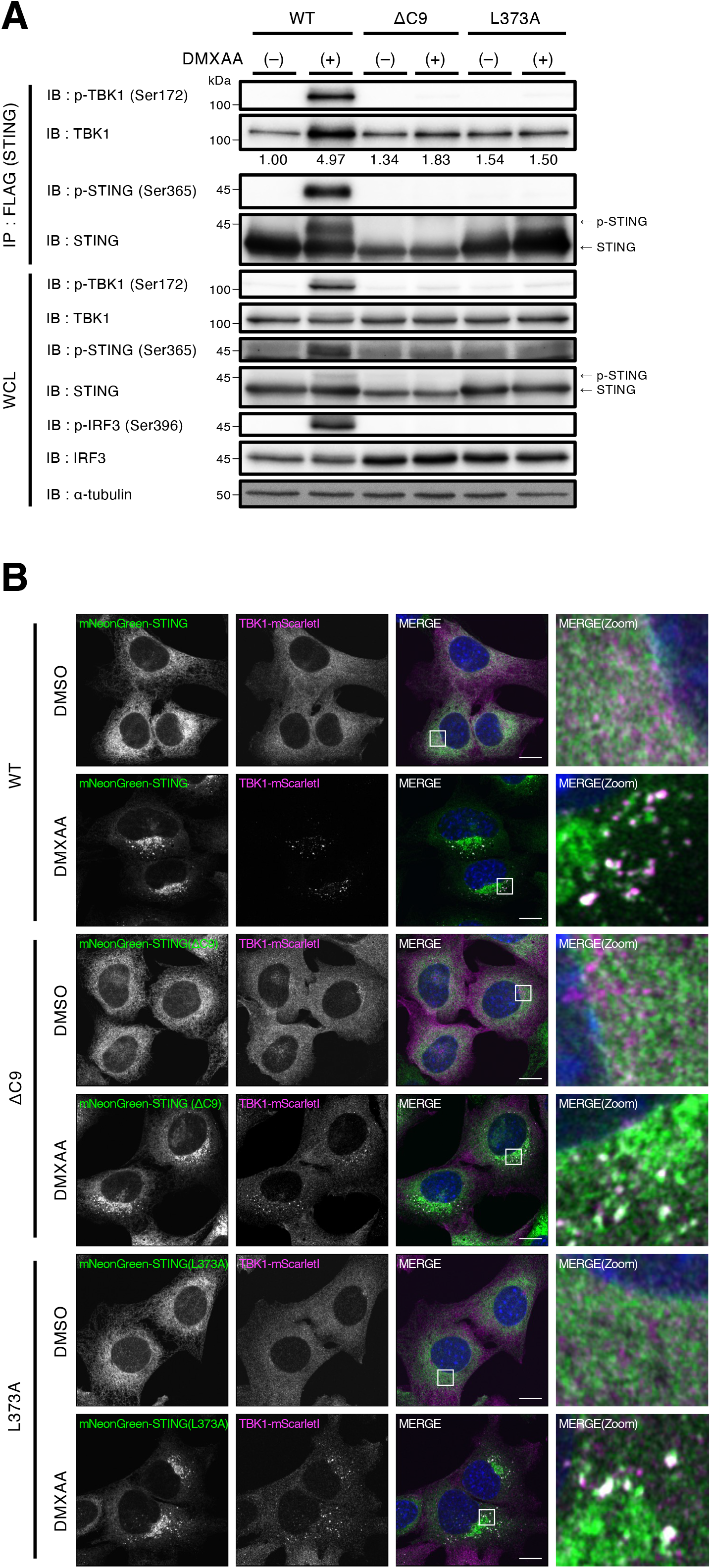
PLPLR[T/S]D motif within the C-terminal tail of STING is required for the binding to TBK1, but is dispensable for the recruitment of TBK1 to the Golgi. (A) FLAG-STING (WT, ΔC9, or L373A) and TBK1-mScarletI expressing ST-DKO MEFs were stimulated with DMXAA (25 μg/mL) for 60 min. Cell lysates were prepared, and FLAG-STING was immunoprecipitated with anti-FLAG antibody. The cell lysates and the immunoprecipitated proteins were analyzed by western blot. The band intensity of co-immunoprecipitated TBK1 was quantified. (B) mNeonGreen-STING (WT, ΔC9, or L373A) and TBK1-mScarletI expressing ST-DKO MEFs were stimulated with DMXAA (25 μg/mL) for 60 min, fixed, permeabilized, and stained with DAPI (blue). Scale bar, 10 μm.

We then examined the recruitment of TBK1-mScarletI to the Golgi after DMXAA stimulation. In MEFs expressing STING (ΔC9) or STING (L373A), as in MEFs expressing WT-STING, TBK1-mScarlet relocated from the cytosol to multiple puncta, albeit with some hazy cytosolic localization left (Figure 3B). At these puncta, TBK1-mScarletI co-localized with the STING variants. These results suggested that the PLPLR[T/S]D motif within the C-terminal tail of STING was dispensable for the recruitment of TBK1 to the Golgi.

### Kinase activity of TBK1 is dispensable for its recruitment to the Golgi

We sought to determine if the kinase activity of TBK1 is involved in its binding to STING or its recruitment to the Golgi. Kinase-dead variant (D135N) or autophosphorylation-deficient variant (S172A) of TBK1, and FLAG-STING were stably expressed in ST-DKO MEFs. After stimulation with DMXAA for 60 min, p-TBK1 (Ser172), p-STING (Ser365), and p-IRF3 (Ser396) emerged in MEFs expressing WT-TBK1, but not in MEFs expressing TBK1 (D135N) or TBK1 (S172A) (Figure 4A), in accordance with the findings that TBK1 is a critical kinase for STING (Ser365) and IRF3 (Ser396) (Liu et al., 2015). We then immunoprecipitated FLAG-STING with anti-FLAG antibody. Co-immunoprecipitated TBK1 increased about 9-fold at 60 min after DMXAA stimulation in MEFs expressing WT-TBK1. In contrast to WT-TBK1, the TBK1 variant (D135N or S172A) showed a reduced binding to STING. These results suggested that kinase activity of TBK1 was essential for its binding to STING. We then examined the recruitment of the TBK1 variants to the Golgi. After DMXAA stimulation, TBK1 (D135N) and TBK1 (S172A), as WT-TBK1, relocated from the cytosol to multiple puncta (Figure 4B). At these puncta, the TBK1 variants co-localized with STING. These results suggested that kinase activity of TBK1 was dispensable for its recruitment to the Golgi.

**Figure 4.**
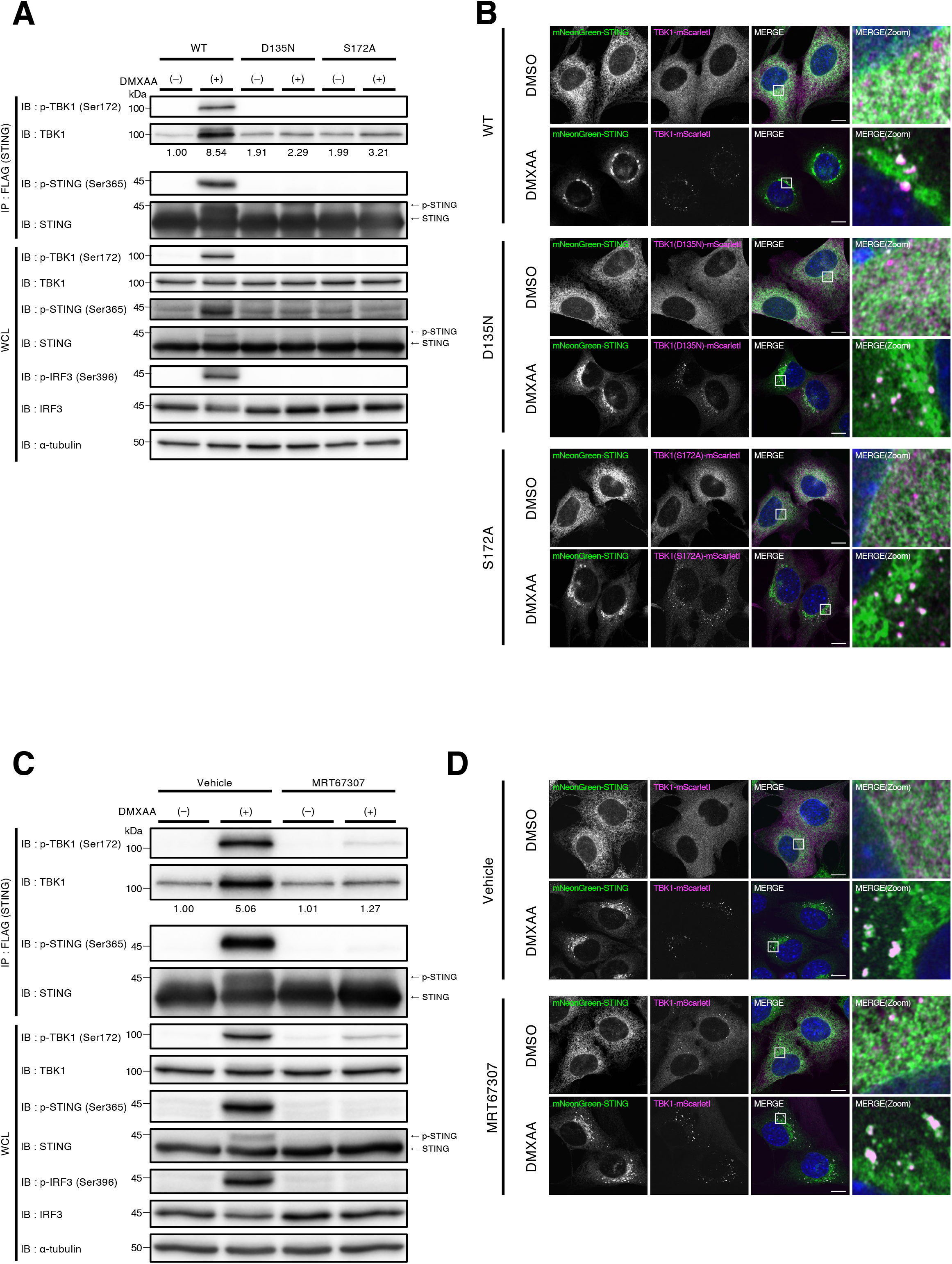
Kinase activity of TBK1 is required for its binding to STING, but is dispensable for TBK1 recruitment to the Golgi. (A) FLAG-STING and TBK1 (WT, D135N, or S172A)-mScarletI expressing ST-DKO MEFs were stimulated with DMXAA (25 μg/mL) for 60 min. Cell lysates were prepared, and FLAG-STING was immunoprecipitated with anti-FLAG antibody. The cell lysates and the immunoprecipitated proteins were analyzed by western blot. The band intensity of co-immunoprecipitated TBK1 was quantified. (B) mNeonGreen-STING and TBK1(WT, D135N, or S172A)-mScarletI expressing ST-DKO MEF were stimulated with DMXAA (25 μg/mL) for 60 min, fixed, permeabilized, and stained with DAPI (blue). (C) FLAG-STING and TBK1-mScarletI expressing ST-DKO MEFs were treated with vehicle or MRT67307 (10 μM) for 2 h, and then stimulated with DMXAA (25 μg/mL) for 60 min. The cell lysates and the immunoprecipitated proteins were analyzed by western blot. The band intensity of co-immunoprecipitated TBK1 was quantified. (D) mNeonGreen-STING and TBK1-mScarletI expressing ST-DKO MEF were treated with vehicle or MRT67307 (10 μM) for 2 h, and then stimulated with DMXAA (25 μg/mL) for 60 min. Cells were fixed, permeabilized, and stained with DAPI (blue). Scale bar, 10 μm.

MRT67307, a TBK1-specific inhibitor (Clark et al., 2011), suppressed the binding of TBK1 to STING (Figure 4C), but not affected the recruitment of TBK1 from the cytosol to multiple puncta (Figure 4D). These results emphasized that kinase activity of TBK1 was essential for its binding to STING, but not for TBK1 recruitment to the Golgi.

## Discussion

In response to the emergence of DNA in the cytosol, STING relocates from the endoplasmic reticulum (ER) to the Golgi and induces the type I interferon response through the activation of cytosolic TBK1. A number of recent studies, essentially by interfering with the COP-II-mediated membrane transport from the ER, indicated that the relocation of STING is required for the activation of STING signalling pathways (Dobbs et al., 2015; Mukai et al., 2016; Ogawa et al., 2018; Gui et al., 2019; Sun et al., 2018; Ran et al., 2019). Moreover, the active form of TBK1, *i.e.*, auto-phosphorylated TBK1, was exclusively confined to subdomains within TGN (Mukai et al., 2016; Mukai et al., 2021). In the present study, using a cell system by which STING and TBK1 are simultaneously monitored with the aid of fluorescent proteins, we found that TBK1 was directly recruited to TGN, not to the other parts of the Golgi (Figures 2 and S1). These results suggested that the Golgi, or TGN in particular, is the site of TBK1 recruitment and activation.

Recent structural studies show that the PLPLR[T/S]D motif within the *C*-terminal tail of STING binds the dimer interface of TBK1 (Zhang et al., 2019; Zhao et al., 2019). Mutations or deletion of amino acid residues in the motif suppress the activation of TBK1, phosphorylation of STING, and the induction of type I interferons, suggesting that the interaction of STING to TBK1 through the *C*-terminal motif is essential for activation of the STING signalling pathways (Zhang et al., 2019; Zhao et al., 2019). We thus examined the contribution of the *C*-terminal motif to TBK1 recruitment to the Golgi in our cell system. Contrary to what we expected, the *C*-terminal motif was dispensable for TBK1 recruitment to the Golgi (Figure 3). However, since the STING variants with mutation or deletion of the *C*-terminal motif lost the interaction with TBK1, the *C*-terminal motif was indeed essential for the interaction of STING and TBK. These results suggested that, after TBK1 was initially recruited onto STING or the Golgi membranes, the *C*-terminal motif strengthened the interaction between STING and TBK, facilitating the formation of stable STING signalling complex. We also found that the recruitment of TBK1 to the Golgi did not require its kinase activity (Figure 4). As with the case in the *C*-terminal motif, the kinase activitiy of TBK1 was essential for the interaction of STING and TBK1 (Figure 4). How the kinase activity of TBK1 contributes to the stable interaction between STING and TBK1 remains to be elucidated.

We previously showed that palmitoylation of STING at the Golgi was required for the induction of type I interferons (Mukai et al., 2016). Treatment of cells with C-176, mouse STING-specific palmitoylation inhibitor (Haag et al., 2018), mostly suppressed the emergence of p-STING and p-IRF3 (Figure S5A). In contrast, p-TBK1 signal did not completely disappear in whole cell lysate or co-immunoprecipitates with FLGA-STING. C-176 did not affect the recruitment of TBK1 to the Golgi (Figure S5B). Thus, palmitoylation of STING, as well as the kinase activity of TBK1 and the *C*-terminal motif of STING, was dispensable for the recruitment of TBK1 to the Golgi. Intriguingly, the number of TBK1 puncta which had an area of more than 0.07 μm^2^ was significantly reduced in cells treated with C-176 (Figure S5C), suggesting that the number of STING/TBK1 contained within single punctum was reduced. The reduction may result in inefficient phosphorylation reactions of STING and IRF3 by TBK1, which was supposed to occur “by induced proximity in trans” across different STING oligomers (Zhao et al., 2019).

## Methods

### Antibodies

Antibodies used in this study were as follows: rabbit anti-phospho-STING (D8F4W, dilution 1:1000 for western blot), rabbit anti-phospho-STING (D1C4T, dilution 1:1000 for immunofluorescence), rabbit anti-phospho-TBK1 (D52C2, dilution 1:1000), rabbit anti-IRF3 (D83B9, dilution 1:1000), rabbit anti-phospho-IRF3 (4D4G, dilution 1:1000) (Cell Signaling Technology); rabbit anti-TBK1 (ab40676, dilution 1:1000) (Abcam); mouse anti-α-tubulin (DM1A, dilution 1:1000) (Sigma-Aldrich); Goat Anti-Rabbit IgG (H + L) Mouse/Human ads-HRP (4050-05, dilution 1:10,000) and Goat Anti-Mouse IgG (H + L) Human ads-HRP (1031-05, dilution 1:10,000) (Southern Biotech); sheep anti-TGN38 (AHP499G, dilution 1:200) (Bio-Rad); mouse anti-GM130 (610823, dilution 1:2000) (BD Biosciences). The antibody against STING was generated by immunizing rabbit with recombinant glutathione S-transferase (GST)–hSTING-C (amino acids 173–379) produced in *E. coli*.

### Reagents

The following reagents were purchased from the manufacturers as noted: DMXAA (14617, Cayman); Nocodazole (13857, Cayman); MRT67307 (19916, Cayman).

### PCR cloning

*N*-terminal mNeonGreen-, EGFP-, or, FLAG-tagged mouse STING (NM_028261) was introduced into pMXs-IPuro. *C*-terminal mScarletI- or mNeonGreen-tagged human TBK1 (NM_013254) was introduced into pMXs-IBla. Variants of STING [ΔC9 (aa 1-369) and L373A)] and TBK1 (D135N and S172A) were generated by site-directed mutagenesis. Mouse GM130 and Rab6 were amplified by PCR with complementary DNA derived from C57BL/6J mouse thymus. The product encoding mouse GM130 was introduced into pMXs-IPuro-mScarletI to generate *N*-terminal mScarletI-tagged GM130. The product encoding mouse Rab6 was introduced into pMXs-IHyg-mScarletI to generate *N*-terminal mScarletI-tagged Rab6.

### Cell culture

MEFs were obtained from embryos of WT or *Sting*^−/−^ mice at E13.5 and immortalized with SV40 Large T antigen. MEFs were cultured in DMEM supplemented with 10% fetal bovine serum (FBS) and penicillin/streptomycin/glutamine (PSG) in a 5 % CO_2_ incubator.

MEFs that stably express tagged proteins were established using retrovirus. Plat-E cells were transfected with pMXs vectors, and the medium that contains the retrovirus was collected. MEFs were incubated with the medium and then selected with puromycin (2 μg/mL), blasticidin (5 μg/mL), or hygromycin (400 μg/mL) for several days.

### Generation of STING- and TBK1-double knockout cells by CRISPR-Cas9

Single-guide RNA (sgRNA) was designed to target mouse TBK1 genomic loci. The sgRNA (sense: 5’-caccgGAGGAGCCGTCCAATGCGTA-3’, antisense: 5’-aaacTACGCATTGGACGGCTCCTCc) was cloned into pSpCas9 (BB)-2A-mCherry created by PX459 V2.0 (Addgene #62988). This plasmid was then transfected into immortalized STING-knockout MEFs (Mukai et al., 2016) with PEI MAX (24765-1, Polyscience). Twenty-four hours after transfection, mCherry expressing cells were sorted by flow cytometry (SH800, SONY). Single colonies were then isolated by limited dilution cloning and the expression of TBK1 in each clone was analyzed by western blot.

### Immunocytochemistry

Cells were fixed with 4% paraformaldehyde (PFA) in PBS at room temperature for 15 min, permeabilized with 0.1% Triton X-100 in PBS at room temperature for 5 min. After blocking with 3 % BSA in PBS, cells were incubated with primary antibodies. After washing with PBS three times, cells were then incubated with the secondary antibody at room temperature for 60 min, washed, and mounted with ProLong™ Glass Antifade Mountant (P36982, Thermo Fisher Scientific).

### Fixed-cell imaging

Cells were seeded on coverslips (13 mm No.1S, MATSUNAMI) the day before fixation. Confocal microscopy was performed using a LSM880 with Airyscan (Zeiss) with a 63 × 1.4 Plan-Apochromat oil immersion lens or 100 × 1.46 alpha-Plan-Apochromat oil immersion lens. Images were analyzed and processed with Zeiss ZEN 2.3 SP1 FP3 (black, 64 bit) (ver. 14.0.21.201) and Fiji (ver. 2.0.0-rc-69/1.52p).

### Immunoprecipitation

Cells were washed with ice-cold PBS and lysed with IP buffer [50 mM HEPES-NaOH (pH 7.2), 150 mM NaCl, 5 mM EDTA, 1% Triton X-100, protease inhibitors (Protease Inhibitor Cocktail for Use with Mammalian Cell and Tissue Extracts, 25955-11, nacalai tesque), and phosphatase inhibitors (8 mM NaF, 12 mM beta-glycerophosphate, 1 mM Na_3_VO_4_, 1.2 mM Na_2_MoO_4_, 5 μM cantharidin, and 2 mM imidazole)]. The cell lysates were centrifugated at 20,000 x g for 15 min at 4 °C, and the resultant supernatants were pre-cleared at 4 °C for 30 min. The lysates were then incubated for 60 min at 4 °C with anti-FLAG M2 Affinity Gel (A2220, Sigma-Aldrich). The beads were washed three times with immunoprecipitation wash buffer [50 mM HEPES-NaOH (pH 7.2), 150 mM NaCl, 0.1 % Triton X-100], eluted with 2 x Laemmli sample buffer, and boiled for 10 min at 95 °C. The immunoprecipitated proteins were separated with SDS-PAGE and transferred to PVDF membrane, then analyzed by western blotting.

### Quantification of the number of TBK1 puncta in the Golgi

Images of TBK1-mNeonGreen and mScarletI-GM130 were thresholded using Yen’s method with Fiji. TBK1- and GM130-positive regions of interest were extracted by multiplying the binarized TBK1 image by the binarized GM130 image. TBK1- and GM130-positive regions of interest were defined using the “Analyze Particles” menu from Fiji on the binary thresholded image. The same processing was applied to the images of TBK1-mNeonGreen and mScarletI-Rab6.

### Quantification of the size of TBK1 puncta

Images of TBK1-mNeonGreen were thresholded using Yen’s method with Fiji. TBK1-positive regions of interest were defined using the “Analyze Particles” menu from Fiji on the binary thresholded image.

### Statistical analysis

Statistical significance was determined with two-tailed Student’s t-test. Data shown are representative of three independent experiments.

## Acknowledgements

We thank Ko Sugawara (Institut de Génomique Fonctionnelle de Lyon, France) for advice on the method of image quantification. This work was supported by JSPS KAKENHI Grant Numbers JP19H00974 (T.T.), JP17H06164 (K.M.), JP17H06418 (K.M.), JP20H03202 (K.M.), JP20H05307 (K.M.), and JP19J21426 (Y.K.); AMED-PRIME (17939604) (T.T.); JST Center of Innovation program from Japan (JPMJCE1303) (K.M.); the Subsidy for Interdisciplinary Study and Research concerning COVID-19 (Mitsubishi Foundation) (T.T.), Takeda Science Foundation (K.M.), Daiichi Sankyo Foundation of Life Science (K.M.), the Cell Science Research Foundation (K.M.), Grant for Basic Science Research Projects from the Sumitomo Foundation (K.M.), Koyanagi-Foundation (K.M.). C-176 was generously provided by Carna Biosciences, Inc.

## Author contribution

H.K. designed and performed the experiments, analyzed the data, interpreted the results, and wrote the paper; Y.K. designed the experiments; K.M. designed the experiments, analyzed the data, interpreted the results, and wrote the paper and; T.T. designed the experiments, interpreted the results, and wrote the paper.

## Competing interests

The authors declare no competing interests.

---

**Figure S1. Representative images related to Figure 2**

Five mini-Golgis indicated by arrowheads in the cell images at the right column were analyzed. The *cis*- and *trans*-regions of the mini-Golgis were outlined in the image. Scale bars, 10 μm.

**Figure S2. Phosphorylated STING localizes to a subdomain of the TGN after stimulation**

(**A**) EGFP-STING expressing STING-knockout MEFs were treated with nocodazole (2.5 μM) for 30 min followed by stimulation with DMXAA (25 μg/mL) for 30 min. Cells were fixed, permeabilized, and stained for GM130 (cyan), TGN38 (magenta), and p-STING (S365) (Red). Scale bars, 10 μm. (**B**) One mini-Golgi indicated by arrowhead in (A) was magnified. The *cis*- and *trans*-regions of the mini-Golgi were outlined in the images at the bottom row. Scale bar, 1 μm. (**C**) Fluorescence intensity profile along the arrows in (B) is shown.

**Figure S3. Co-localization analysis of CGN and TGN proteins with Airyscan super-resolution microscopy**

(**A**) mScarletI-GM130 expressing MEFs were fixed, permeabilized, and stained for endogenous TGN38 (a TGN protein). (**B**) Fluorescence intensity profile along the arrow in (A) is shown. (**C**) mScarletI-Rab6 expressing MEFs were fixed, permeabilized, and stained for endogenous GM130 (a CGN protein). (**D**) Fluorescence intensity profile along the arrow in (C) is shown. Scale bar, 10 μm.

**Figure S4. TBK1 is recruited from the cytosol to TGN**

(**A** to **H**) TBK1-knockout MEFs were reconstituted with TBK1-mNeonGreen using retrovirus. mScarletI-GM130 (a CGN protein) (A to D) or mScarletI-Rab6 (a TGN protein) (E to H) was stably expressed in the reconstituted MEFs. Cells were then stimulated with DMXAA (25 μg/mL) for 20 min and fixed.

(**A**) The fluorescence images of TBK1 and GM130.

(**B**) Binarized images of (A).

(**C**) Outlines of the binarized images of (B) were obtained, and the outlined regions of interest were merged with the images of (A).

(**D**) TBK1-positive puncta inside or outside CGN (GM130) were determined using the data in (C), and outlined with green or yellow, respectively. The outlined regions of interest were then merged with the image of TBK1-mNeonGreen in (A). The number of TBK1-positive puncta inside CGN was counted in (I).

(**E**) The fluorescence images of TBK1 and mScarletI-Rab6.

(**F**) Binarized images of (E).

(**G**) Outlines of the binarized images of (F) were obtained, and the outlined regions of interest were merged with the images of (E).

(**H**) TBK1-positive puncta inside or outside TGN (Rab6) were determined using the data in (G), and outlined with green or yellow, respectively. The outlined regions of interest were then merged with the image of TBK1-mNeonGreen in (E). The number of TBK1-positive puncta inside TGN was counted in (I).

(**I**) The number of TBK1 puncta inside CGN (mScarletI-GM130) in (D) or TGN (mScarletI-Rab6) in

(H) was plotted. Data are from two (n≧18 cells) independent experiments. Statistical significance was determined with two-tailed Student’s *t*-test.

**Figure S5. Inhibition of STING palmitoylation does not affect the recruitment of TBK1 to the Golgi after stimulation**

(**A**) FLAG-STING and TBK1-mScarletI expressing ST-DKO MEFs were treated with vehicle or C-176 (10 μM) for 2 h, and then stimulated with DMXAA (25 μg/mL) for 60 min. The cell lysates and the immunoprecipitated proteins were analyzed by western blot. The band intensity of co-immunoprecipitated TBK1 was quantified. (**B**) mNeonGreen-STING and TBK1-mScarletI expressing ST-DKO MEF were treated with vehicle or C-176 (10 μM) for 2 h, and then stimulated with DMXAA (25 μg/mL) for 60 min. Cells were fixed, permeabilized, and stained with DAPI (blue). Scale bar, 10 μm. (**C**) The number of TBK1 puncta which had an area of more than 0.07 μm^2^ in (B) were counted. Data are from ten cells. Statistical significance was determined with two-tailed Student’s *t*-test.

## Notes

### Competing Interest Statement

The authors have declared no competing interest.

