## Supplementary figures for "Kinase activity of TBK1 is required for its binding to STING, but not for its recruitment to the Golgi"

### Supplementary Figure 1

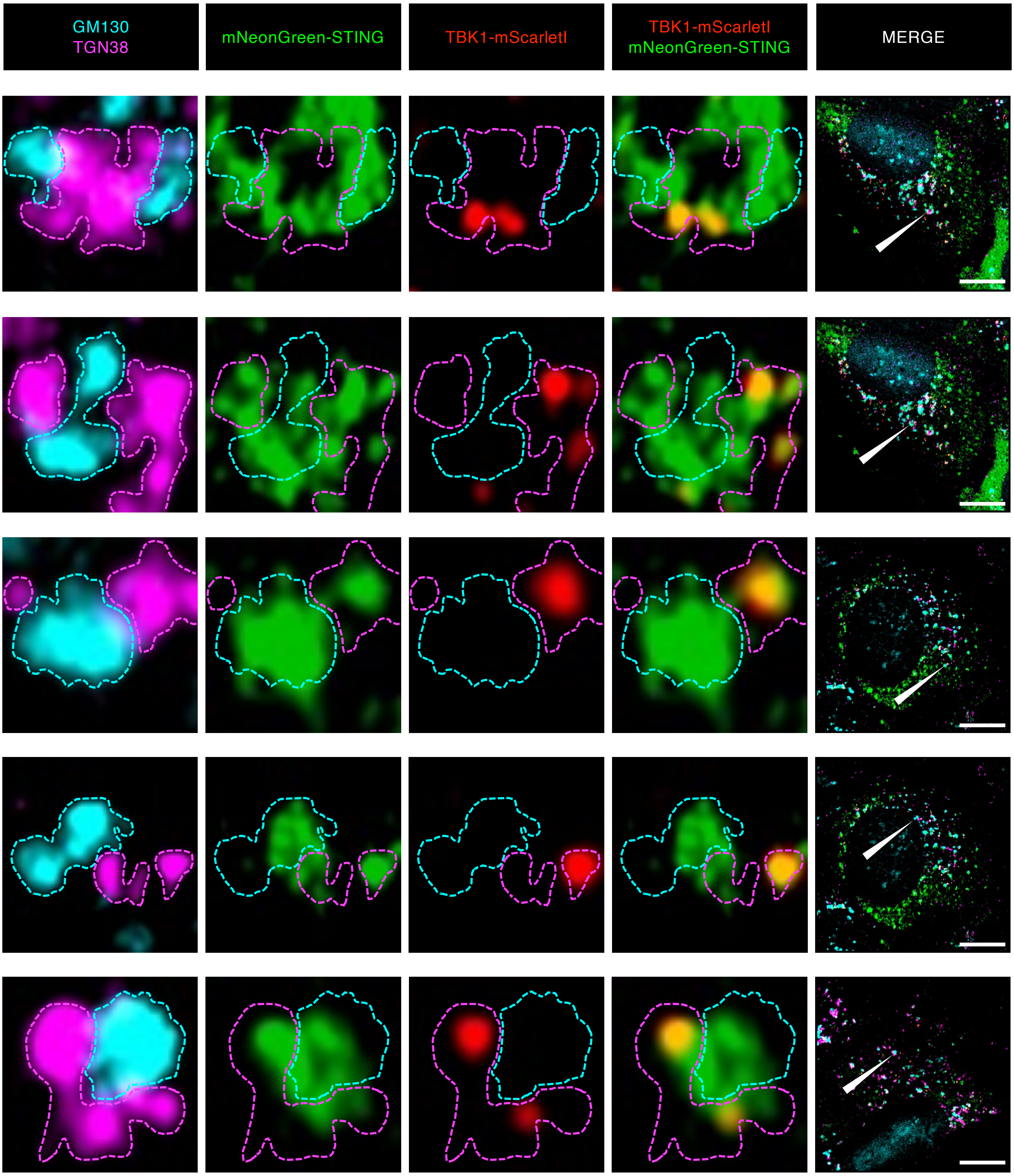


**Figure S1. Representative images related to Figure 2**
Five mini-Golgis indicated by arrowheads in the cell images at the right column were analyzed. The *cis*- and *trans*-regions of the mini-Golgis were outlined in the image. Scale bars, 10 µm.

### Supplementary Figure 2

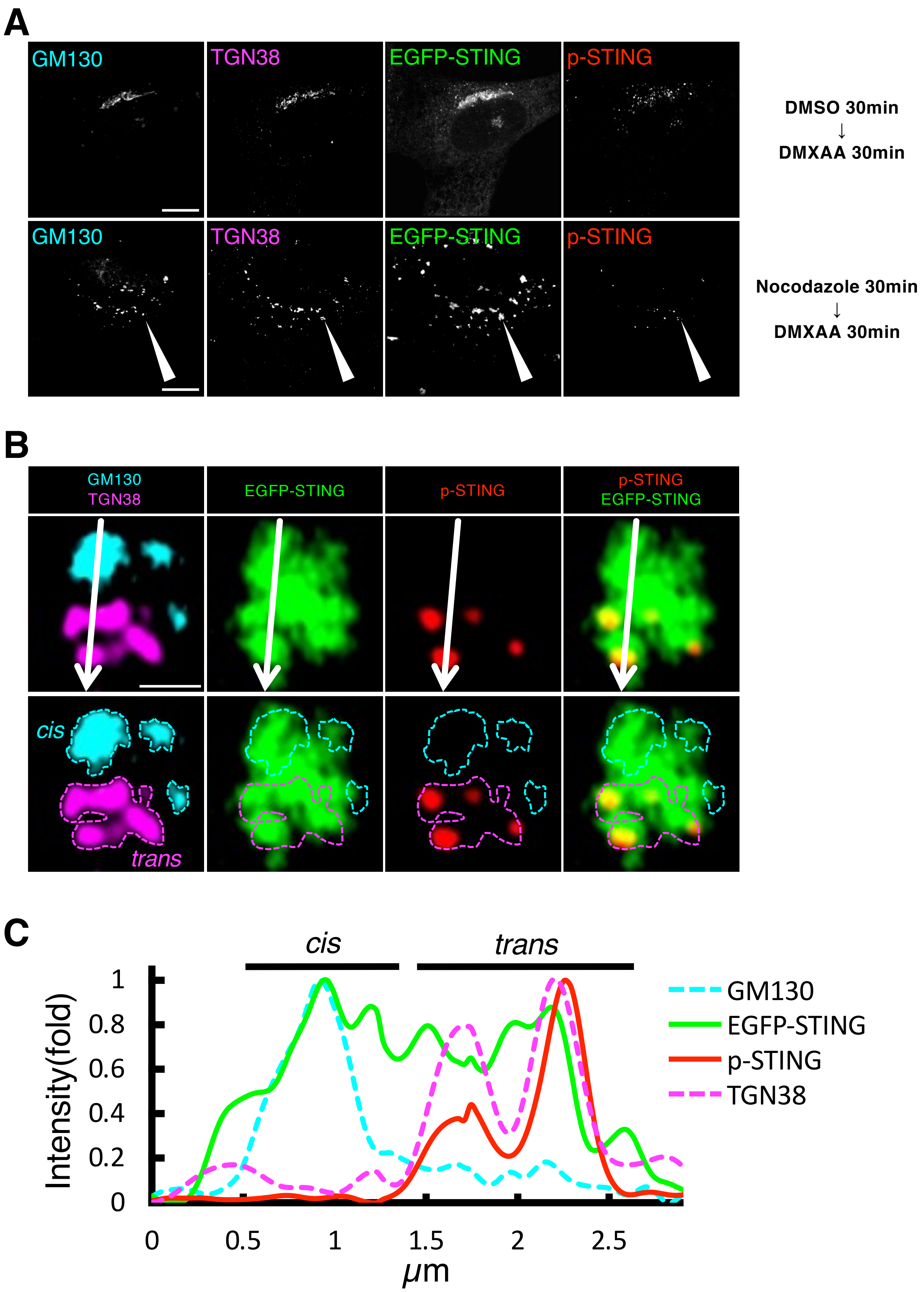


**Figure S2. Phosphorylated STING localizes to a subdomain of the TGN after stimulation**
(**A**) EGFP-STING expressing STING-knockout MEFs were treated with nocodazole (2.5 μM) for 30 min followed by stimulation with DMXAA (25 μg/mL) for 30 min. Cells were fixed, permeabilized, and stained for GM130 (cyan), TGN38 (magenta), and p-STING (S365) (Red). Scale bars, 10 μm. (**B**) One mini-Golgi indicated by arrowhead in (A) was magnified. The *cis*- and *trans*-regions of the mini-Golgi were outlined in the images at the bottom row. Scale bar, 1 μm. (**C**) Fluorescence intensity profile along the arrows in (B) is shown.

### Supplementary Figure 3

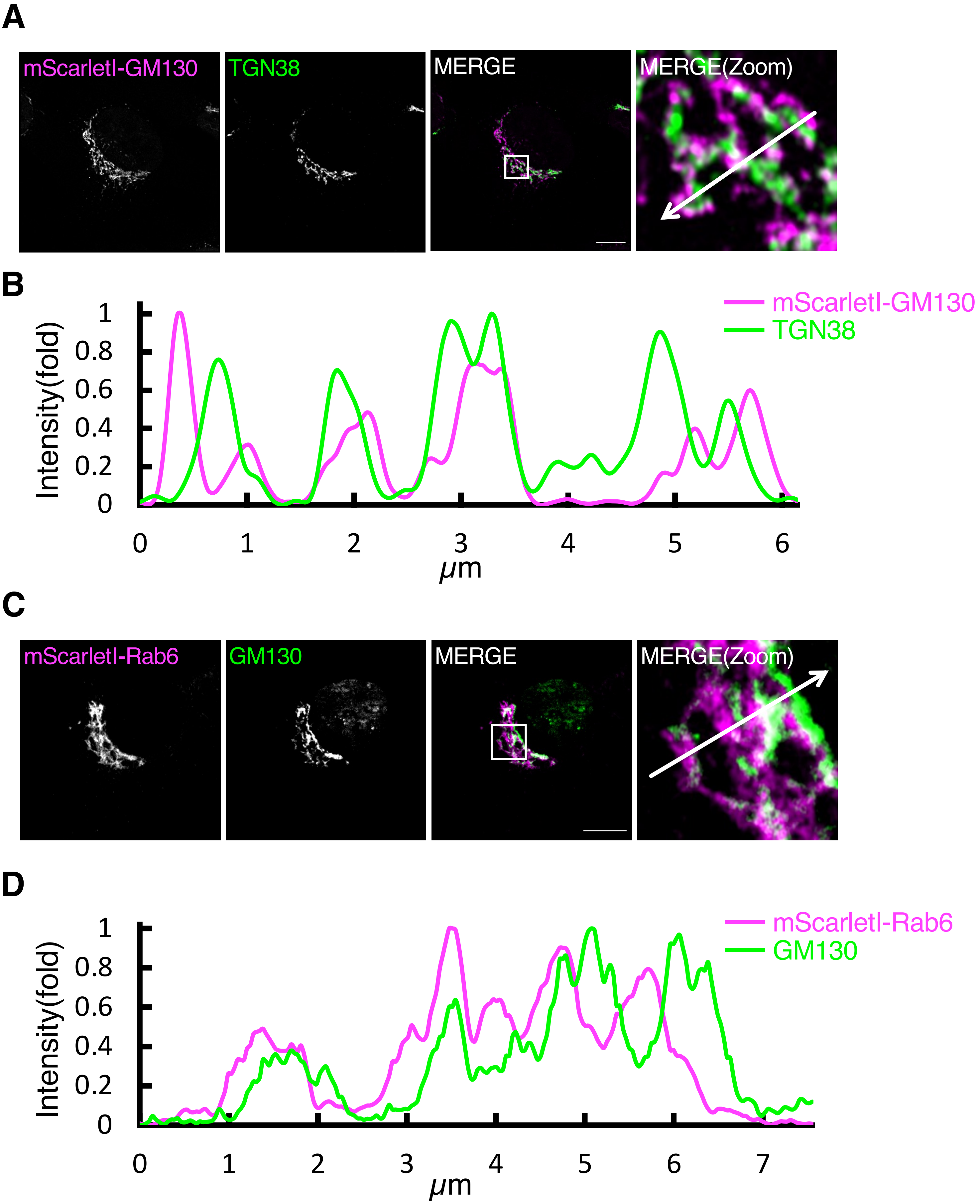


**Figure S3. Co-localization analysis of CGN and TGN proteins with Airyscan super-resolution microscopy**

(**A**) mScarletI-GM130 expressing MEFs were fixed, permeabilized, and stained for endogenous TGN38 (a TGN protein). (**B**) Fluorescence intensity profile along the arrow in (A) is shown. (**C**) mScarletI-Rab6 expressing MEFs were fixed, permeabilized, and stained for endogenous GM130 (a CGN protein). (**D**) Fluorescence intensity profile along the arrow in (C) is shown. Scale bar, 10 μm.

### Supplementary Figure 4

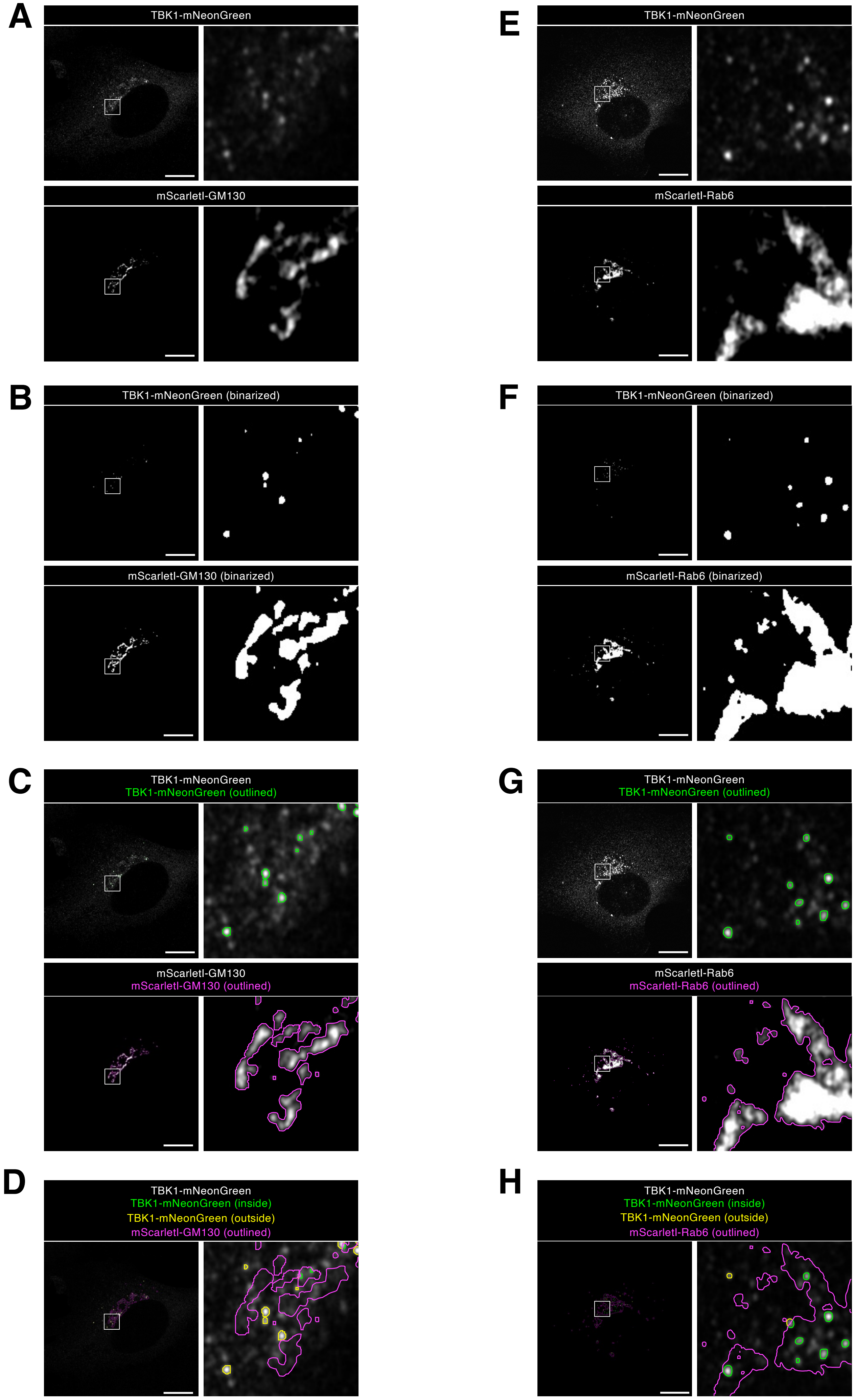

### Supplementary Figure 4

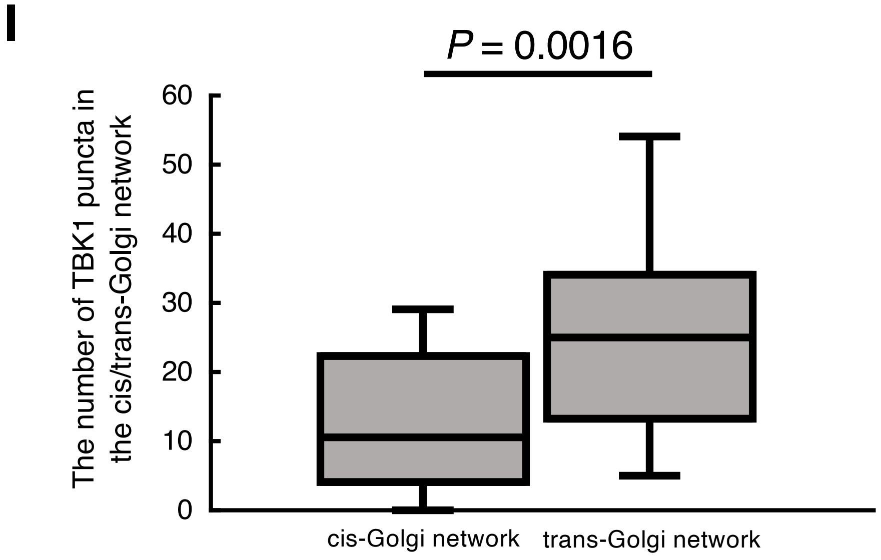


**Figure S4. TBK1 is recruited from the cytosol to TGN**

(**A** to **H**) TBK1-knockout MEFs were reconstituted with TBK1-mNeonGreen using retrovirus. mScarletI-GM130 (a CGN protein) (A to D) or mScarletI-Rab6 (a TGN protein) (E to H) was stably expressed in the reconstituted MEFs. Cells were then stimulated with DMXAA (25 µg/mL) for 20 min and fixed.

(**A**) The fluorescence images of TBK1 and GM130.

(**B**) Binarized images of (A).

(**C**) Outlines of the binarized images of (B) were obtained, and the outlined regions of interest were merged with the images of (A).

(**D**) TBK1-positive puncta inside or outside CGN (GM130) were determined using the data in (C), and outlined with green or yellow, respectively. The outlined regions of interest were then merged with the image of TBK1-mNeonGreen in (A). The number of TBK1-positive puncta inside CGN was counted in (I).

(**E**) The fluorescence images of TBK1 and mScarletI-Rab6.

(**F**) Binarized images of (E).

(**G**) Outlines of the binarized images of (F) were obtained, and the outlined regions of interest were merged with the images of (E).

(**H**) TBK1-positive puncta inside or outside TGN (Rab6) were determined using the data in (G), and outlined with green or yellow, respectively. The outlined regions of interest were then merged with the image of TBK1-mNeonGreen in (E). The number of TBK1-positive puncta inside TGN was counted in (I).

(**I**) The number of TBK1 puncta inside CGN (mScarletI-GM130) in (D) or TGN (mScarletI-Rab6) in (H) was plotted. Data are from two ($n \geqq 18$ cells) independent experiments. Statistical significance was determined with two-tailed Student's *t*-test.

### Supplementary Figure 5

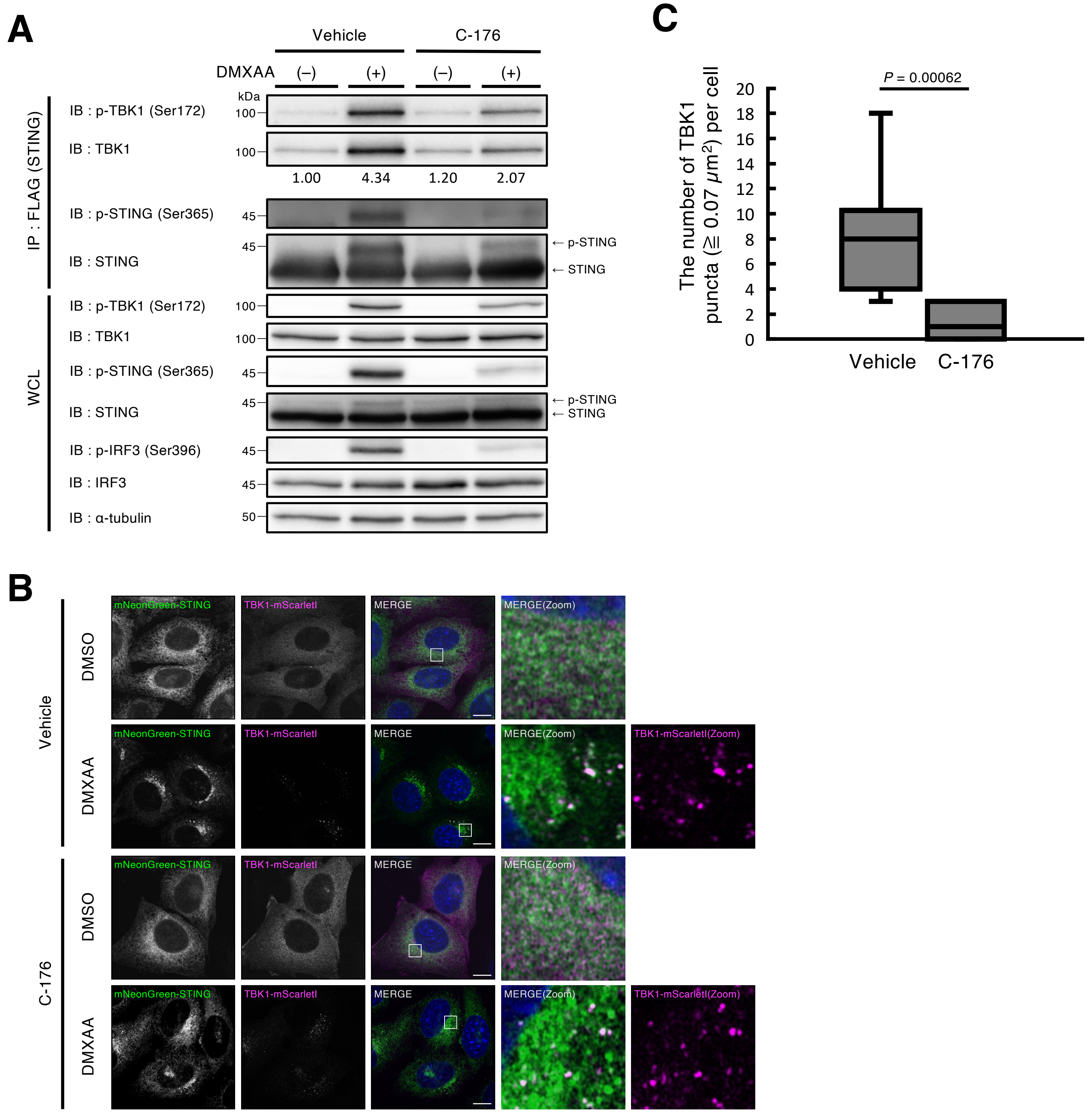


**Figure S5. Inhibition of STING palmitoylation does not affect the recruitment of TBK1 to the Golgi after stimulation**

(**A**) FLAG-STING and TBK1-mScarletI expressing ST-DKO MEFs were treated with vehicle or C-176 (10 µM) for 2 h, and then stimulated with DMXAA (25 µg/mL) for 60 min. The cell lysates and the immunoprecipitated proteins were analyzed by western blot. The band intensity of co-immunoprecipitated TBK1 was quantified. (**B**) mNeonGreen-STING and TBK1-mScarletI expressing ST-DKO MEF were treated with vehicle or C-176 (10 µM) for 2 h, and then stimulated with DMXAA (25 µg/mL) for 60 min. Cells were fixed, permeabilized, and stained with DAPI (blue). Scale bar, 10 µm. (**C**) The number of TBK1 puncta which had an area of more than 0.07 µm$^2$ in (B) were counted. Data are from ten cells. Statistical significance was determined with two-tailed Student's t-test.
